# Quantifying Differential Rhythmicity based on Effect Sizes with LimoRhyde2

**DOI:** 10.1101/2024.05.09.593377

**Authors:** Dora Obodo, Amir Asiaee

**Affiliations:** Department of Biostatistics, St. Jude Children’s Research Hospital, Memphis, TN, USA; Department of Biostatistics, Vanderbilt University Medical Center, Nashville, TN, USA

## Abstract

Current methods for assessing differential rhythmicity in genomic data focus on hypothesis testing and model selection, often assuming sinusoidal rhythms. A more appropriate approach is to estimate differences in rhythmic properties between two or more conditions using effect sizes. To address this gap, we extend LimoRhyde2, a method for quantifying rhythm-related effect sizes and their uncertainty in genome-scale data, to enable differential rhythmicity analyses. Through extensive testing, we validate the method for differential rhythmicity analysis and showcase how it improves biological interpretation for circadian systems biology.

## INTRODUCTION

Genomic analyses have grown increasingly complex as researchers seek to understand circadian systems and their relevance to human health. Circadian genomic experiments typically employ time-series designs spanning one or more 24-hour periods, which help to determine whether genomic features exhibit rhythmic transcription (Hughes et al., 2010). These experiments often involve multiple time point replicates, various tissue types, and different conditions or treatment groups (Hughes et al., 2017). Notably, genome-scale analyses have revealed mechanisms of inter-cellular communication in the circadian response to feeding(Guan et al., 2020), highlighted drug targets for circadian medicine (Ruben et al., 2018), and identified rhythmic transcriptional reprogramming in response to lifestyle-based diseases (Eckel-Mahan et al., 2013). These experiments are imperative to unravel distinct circadian mechanisms and their associated biological processes.

For these experiments, rhythmicity and differential rhythmicity detection enable circadian rhythm analyses. While rhythmicity detection identifies genomic rhythms in experiments where the variable of interest is time, differential rhythmicity helps to analyze how gene rhythms respond to environmental and genetic changes. More specifically, differential rhythmicity includes changes to rhythm parameters such as amplitude, phase, or both. In addition, changes to baseline expression, or mesor, of a rhythmic gene result in differential expression (Parsons et al., 2020, Singer and Hughey, 2019). Given their distinct goals, statistical methods for detecting rhythmicity in one condition are ill-suited for detecting differential rhythmicity between conditions, and their inappropriate use can lead to misinterpretation (Ness-Cohn et al., 2021, Pelikan et al., 2022).

Current methods primarily use statistical testing to detect differential rhythmicity and some have integrated their use with standard genomic analysis pipelines (Weger et al., 2021). These methods can be summarized as either performing model selection (Pelikan et al., 2022, Weger et al., 2021) or hypothesis testing (Ding et al., 2021, Parsons et al., 2020, Singer and Hughey, 2019, Thaben and Westermark, 2016, Xue et al., 2023). Broadly, model selection methods aim to identify sets of functions that best describe changes between two conditions, while hypothesis testing evaluates the presence or absence of differential rhythmicity. However, most methods assume that rhythms are sinusoidal, an assumption that previous circadian transcriptome analyses indicate is marginally untrue (Thaben and Westermark, 2014, Zhang et al., 2014). Similarly, reporting p-values and statistical significance alone has several limitations (Obodo et al., 2023). While model selection circumvents some limitations of hypothesis testing, the number of models increases exponentially with the number of conditions, making it hard to scale, and can complicate interpretations (Obodo et al., 2023, Pelikan et al., 2022). Therefore, although methods to assess differential rhythmicity are increasingly popular, they still struggle to reliably quantify differential rhythmicity in genomic analyses with multiple conditions. A more appropriate approach to differential rhythmicity analysis may be to estimate effect sizes for differences in rhythm properties individually. LimoRhyde2 is a method designed to quantify rhythm-related effect sizes and their uncertainty in genome-scale data with one or more conditions (Obodo et al., 2023). However, the method has not been applied to differential rhythmicity analysis. Outside of biological rhythms research, the broader field of genomics has produced multiple well-validated methods for accurately estimating effects(Love et al., 2014, Urbut et al., 2019, Zhu et al., 2019), which has become an important part of differential expression analysis. Unfortunately, limited options exist for estimating differential effects related to biological rhythms. To address this challenge, we extend LimoRhyde2’s statistical framework to quantify differential rhythmicity in datasets with multiple conditions. Here, we conduct extensive testing using artificial experiments to validate the method in various settings and showcase how LimoRhyde2 reliably quantifies differential rhythmicity to improve biological interpretation when investigating circadian systems biology.

## MATERIALS AND METHODS

LimoRhyde2 is available as a comprehensive R package https://limorhyde2.hugheylab.org. Building on the foundation laid for quantifying rhythmicity within single conditions (Obodo et al., 2023), we demonstrate LimoRhyde2’s approach to quantifying differential rhythmicity in multi-conditional data.

### LimoRhyde2 Model for Differential Rhythmicity

#### Spline-based Linear Model

By default LimoRhyde2 utilizes periodic cubic spline linear functions (or optionally, sine and cosine functions), to model circadian time. The algorithm then computes model coefficients via multivariate adaptive shrinkage (**Mash**) which moderates the raw coefficients using the empirical Bayes method.

For multi-condition data, the model is designed to capture the dynamic interactions between temporal patterns and experimental conditions in gene expression data. It is mathematically represented as:

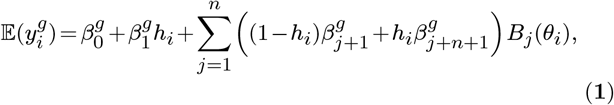

where:

- 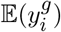: Expected log-transformed measured expression of gene *g* in sample *i*.
- 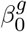: Baseline expression level for gene *g*.
- 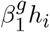: Adjustment to baseline based on the condition *h*_*i*_ (0 for wild-type, 1 for the under-studied condition, e.g., knockout).
- *B*_*j*_ (*θ*_*i*_): The *j*th periodic spline functions modeling the time-related effects, with 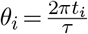translating time *t*_*i*_ into a phase within period *τ*.
- 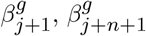: Coefficients for the *j*-th spline function under each condition.

This structure allows the model to effectively describe how gene expressions vary not only over time but also across different experimental conditions, capturing potential shifts in rhythmic patterns due to experimental interventions.

#### Calculating Rhythm and Differential Rhythm Statistics

Using the moderated or optionally the raw coefficients from the model, LimoRhyde2 computes various rhythm statistics for each genomic feature across all conditions. These statistics are derived from the properties of the fitted curve within the interval [0,*τ*]. Key metrics, defined in Table 1, include:

**Table 1.** Definitions of Rhythmic Statistics of function *f* (*t*) with period *τ*.

| Statistic | Definition |
| --- | --- |
| Mesor | Average level, $\frac{1}{\tau} \int_0^\tau f(t) dt$ |
| Peak | Highest value, $\max f(t)$ |
| Trough | Lowest value, $\min f(t)$ |
| Amplitude | Peak to trough distance, $\max f(t) - \min f(t)$ |
| Peak Phase | Time $t_p$ of peak, $\max f(t)$ occurrence |
| Trough Phase | Time $t_t$ of trough, $\min f(t)$ occurrence |

- **Mean Values**: Calculates averages of metrics like mesor and amplitude across conditions, reflecting overall rhythmic expression.
- **Differential Statistics**: Assesses changes between conditions by computing differences in mesor, amplitude, and circular phase differences, highlighting condition-induced rhythmic variations.

These statistics help characterize the dynamics of biological rhythms, providing insights into the timing, magnitude, and consistency of oscillatory patterns under different experimental conditions. For a detailed explanation of each statistic, see Table 1.

LimoRhyde2 quantifies uncertainty in the fitted curves by sampling from the posterior distribution computed by Mash to moderate model coefficients. Using these sampled coefficients, LimoRhyde2 computes credible intervals—the Bayesian equivalent of a confidence interval—for both fitted curves and associated rhythm (differential) statistics, thereby enhancing the reliability of the rhythmic assessments.

### Processing Circadian Transcriptome Data from Mice

To analyze microarray data from mouse liver (GSE11923), we used the seeker R package (Schoenbachler and Hughey, 2022) to download sample metadata and raw Affymetrix files from the NCBI GEO, map probes to Entrez Gene IDs (Dai et al., 2005) and perform robust multi-array average (RMA) normalization (Irizarry et al., 2003), yielding log2-transformed expression values.

For RNA-seq datasets from liver (GSE143524) and SCN (GSE72095), we used the seeker package for metadata retrieval and to download and process raw reads. The preprocessing pipeline included adapter and quality trimming with Trim Galore (Krueger et al., 2021), quality control via FastQC (Andrews, 2023) and MultiQC (Ewels et al., 2016), and gene-level count and abundance quantification using salmon (Patro et al., 2017) and tximport (Soneson et al., 2016), with gene annotations based on Ensembl Gene IDs. The transcriptome index required for salmon was constructed using refgenie (Stolarczyk et al., 2020).

### Quantifying Differential Rhythmicity using LimoRhyde2

To quantify differential rhythmicity for microarray data GSE11923, we ran LimoRhyde2 with four internal knots per spline and a fixed period of 24 hours. We opted for the `limma-trend` approach, which is tailored for microarray data analysis. For the RNA-seq datasets GSE143524 and GSE72095, we applied a filter retaining only genes with counts per million (CPM) of at least 0.5 in a minimum of 75% of samples, regardless of condition or timepoint. To mitigate the impact of zero counts on log-transformed CPM values and prevent overestimated effect sizes, counts of zero were adjusted to the smallest non-zero count observed across all samples for the respective gene. Subsequently, we ran LimoRhyde2 on each dataset with three knots—reflecting the coarser temporal resolution of sample collection—and a fixed 24-hr period. We employed `limma-voom` for linear modeling of log-transformed counts. For relevant analyses, 90% equal-tailed credible intervals were computed from 200 posterior samples, providing a Bayesian estimate of interval confidence.

### Testing Accuracy of LimoRhyde2 Differential Rhythmicity Analysis

#### Selecting Rhythmic Genes

Differential rhythmicity is predicated on the assumption of rhythmic gene expression in at least one condition. To rigorously test LimoRhyde2’s accuracy in quantifying differential rhythmicity, we first tested for gene rhythmicity using p-value-based methods: LimoRhyde (Singer and Hughey, 2019), RAIN (Thaben and Westermark, 2014) and JTK Cycle (Hughes et al., 2010). We then used Stouffer’s method to integrate the three sets of p-values and adjusted the resulting p-values using the Benjamini-Hochberg method to control the false discovery rate (Benjamini and Hochberg, 1995, Yoon et al., 2021). We selected genes with adjusted p-values (q-values) less than 0.01. With these identified genes, we constructed artificial scenarios of differential rhythmicity from real circadian transcriptome data. We then applied LimoRhyde2 to evaluate its performance against the artificial, yet known ground truths.

#### Generating Conditions with Artificial Differential Rhythmicity

After pre-selecting rhythmic genes in a single-condition dataset, we generated artificial differential rhythmicity by creating two conditions: ‘*original*’ and ‘*scaled*.’ The **original** condition was the initial dataset, 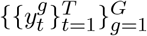 with no sample modifications, where *T* is the total time steps and *G* is the total selected genes. We created the **scaled** condition by scaling sample measurements for each gene to deliberately change the amplitude (or phase), thereby instigating artificial differential rhythmicity which we show with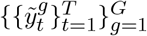.

To scale amplitudes, we randomly assigned scale factors to each gene from a truncated Gaussian distribution: *s*^*g*^ *∼ N* (0.5,0.25), limited to the interval [0,2]. The scale factors were then applied to the original data to generate the scaled data according to the following transformation:

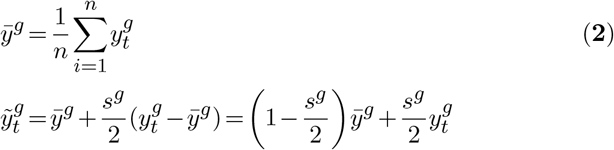

Here, 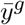 represents the mean expression of gene *g, s*^*g*^ is the gene-specific scale factor, 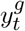 denotes the expression level of gene *g* in time (sample) *t*, and 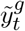 is the adjusted expression for gene *g* in sample *t* after scaling the deviation from the mean, 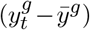, by 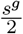 which will scale the amplitude by *s*^*g*^ We performed differential rhythmicity analysis on the resulting data using LimoRhyde2.

#### Evaluating LimoRhyde2 Accuracy

Our objective is to assess the accuracy of LimoRhyde2 in estimating differences in amplitude between two conditions using artificially scaled data. We aim to evaluate both the estimated Δamplitudes and the scale factors.

First, we define the ground truth Δamplitude for the scaled condition based on the scale factor applied. We estimate the original amplitude 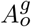 of gene *g* using LimoRhyde2 and assume the ground truth amplitude for the scaled data as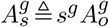, where *s*_*g*_ is the applied scale factor. This defines the ground truth differential amplitude as 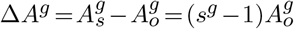. The LimoRhyde2 estimated differential amplitude is then 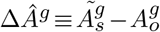, where 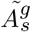 is the amplitude estimated from the scaled data.

We also calculate the estimated scale factor from LimoRhyde2 as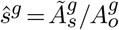, where 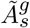 and 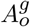 are the estimated scaled and original amplitudes, respectively. To quantify LimoRhyde2’s accuracy in detecting introduced differential rhythmicity, we use the mean absolute error (MAE). The MAE is calculated as follows for the differential amplitudes and scale factors across *G* genes:

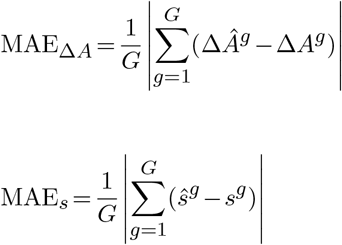

#### Evaluating LimoRhyde2 Under Bootstrap-Induced Variability

We employed bootstrap resampling to quantify the distribution of the Mean Absolute Error (**MAE**) in the estimation of Δamplitude and scaling factors. Utilizing the biological replicates available at each time point *t*, we generated bootstrap time series data. Initially, as described previously, we created ‘original’ and ‘scaled’ conditions with scaling factors drawn from a normal distribution *N* (1,0.35). For each gene in each condition, we randomly selected one of the three replicates at each time point to assemble 500 bootstrap time series pairs. Differential rhythmicity analysis was then conducted using LimoRhyde2 on each paired dataset.

#### Generating Conditions with Null Differential Rhythmicity

To generate null differential rhythmicity, we assigned samples from even-numbered time points to an ‘even’ condition and those from odd-numbered time points to an ‘odd’ condition in a single-condition dataset. Differential rhythmicity analysis was then performed using LimoRhyde2. We adapted this scenario from the methodology used in Pelikan et al (Pelikan et al., 2022).

## RESULTS

To demonstrate how LimoRhyde2 quantifies differential rhythmicity, we applied it to several transcriptome data (Table 2). LimoRhyde2 quantifies differential rhythmicity by fitting a periodic spline-based linear model to each gene in each condition, (raw fits) and moderating the raw fits using Mash (producing posterior fits). It then calculates and outputs each gene’s rhythm statistics (based on the raw and posterior fits), and differential rhythm statistics (between pairs of conditions) (Fig. 1).

**Table 2.**
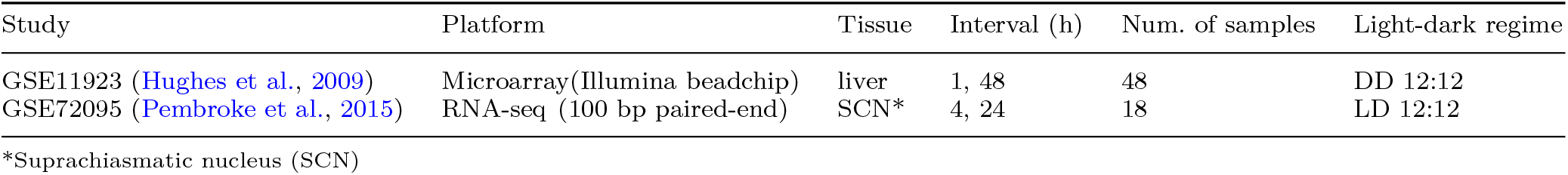
Details of circadian transcriptome datasets used for validation.

| Study | Platform | Tissue | Interval (h) | Num. of samples | Light-dark regime |
| --- | --- | --- | --- | --- | --- |
| GSE11923 (Hughes et al., 2009) | Microarray (Illumina beadchip) | liver | 1, 48 | 48 | DD 12:12 |
| GSE72095 (Pembroke et al., 2015) | RNA-seq (100 bp paired-end) | SCN* | 4, 24 | 18 | LD 12:12 |

\*Suprachiasmatic nucleus (SCN)

**Figure 1.**
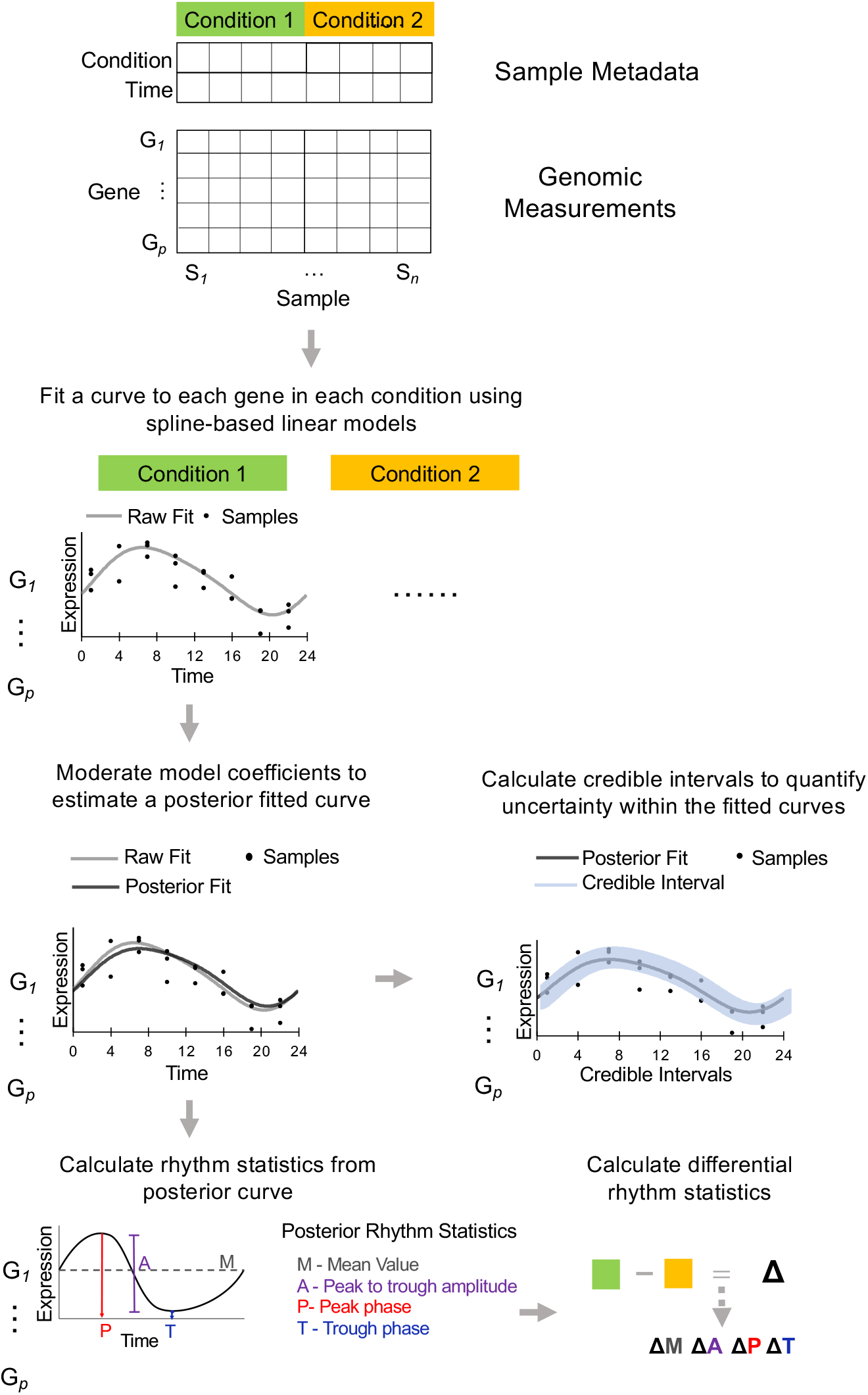
LimoRhyde2 procedure for quantifying differential rhythmicity in genomic data. Given a gene (row) by sample (column) matrix of measurements possibly from multiple conditions (color indicates condition), (1) LimoRhyde2 fits a curve (dashed gray lines) to each gene’s measurements in each condition. (2) To adjust for noise and uncertainty in the fit, LimoRhyde2 moderates the model coefficients, yielding a posterior fit (solid black lines) for each gene in each condition. (3) LimoRhyde2 optionally calculates 90% credible intervals (blue shading) for the posterior fitted curve. (4) Using the posterior fits, LimoRhyde2 calculates rhythm statistics (colored text) for each gene in each condition and differential rhythm statistics (Δ) for each gene between each pair of conditions.

To understand how well LimoRhyde2 quantifies differential rhythmicity, we performed accuracy testing by calculating errors between LimoRhyde2 estimates and ground truths in artificial scenarios of differential rhythmicity. Typically, p-value-based methods may evaluate accuracy by estimating false positives and false negatives (false discoveries) when the differential rhythmicity status for a set of features is known (Pelikan et al., 2022). In this context, a well-validated method may limit these false discoveries. Similarly, a well-validated estimation-based method may minimize error between estimated values and known effect sizes for differential rhythm parameters (e.g., Δamplitude, Δphase). This includes null differential rhythmicity, where estimated values are close to 0 when there is no change in rhythmicity between two conditions. Thus, we evaluated LimoRhyde2 estimation error in cases of null and artificial differential rhythmicity.

To artificially derive differential rhythmicity, we used data from Hughes et al, where wild-type mouse liver gene expression was measured via microarrays every hour for 48 hours (single condition) (Hughes et al., 2009). Differential rhythmicity assumes that a gene is rhythmic in at least one condition. Thus, we first identified rhythmic genes using p-value and amplitude criteria, resulting in a shortened dataset with roughly 4,800 genes out of the original 18,243 (Fig. 2A). To create two conditions, we took the unaltered ‘original’ data to be one condition and generated a second ‘scaled’ condition, where resulting gene amplitudes are scaled up or down based on a randomly assigned scale factor (Fig. 2B) (see Methods section for formulas used to scale). We then performed differential rhythmicity analysis on the two conditions using LimoRhyde2 (Fig. 2B).

**Figure 2.**
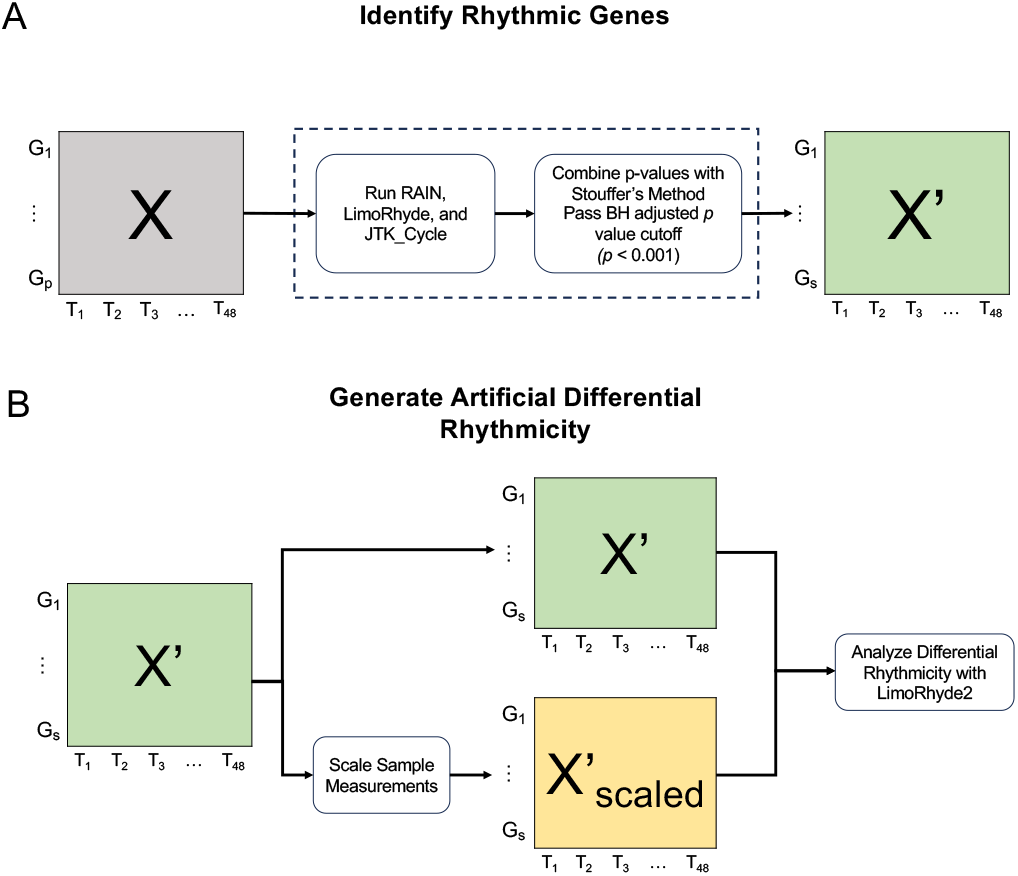
Generating artificial scenarios of differential rhythmicity. **(A)** Given a dataset X (gray square), with genes 1 to p and samples at time points T1 to T48, genes are filtered using LimoRhyde, RAIN, and JTK Cycle algorithms to produce dataset X’ (green squares). **(B)** Two artificial conditions for LimoRhyde2’s differential rhythmicity analysis are created by scaling gene measurements in X’ to produce dataset X’_scaled_ (yellow square).

In the scaled condition, we randomly selected scale factors from a truncated normal distribution bounded between 0 and 2, with a mean of 0.5 (Guan et al., 2020) and s.d. 0.25 (Fig. 3A). These specifications simulate a differential rhythmicity scenario in which most genes will experience a decrease in amplitude. Genes with a scale factor of 1 received no amplitude change, while genes with scale factors less than or greater than 1 had their amplitudes decreased or increased respectively (Fig. 3B). After performing differential rhythmicity analysis, LimoRhyde2 tended to correctly estimate Δamplitude between original and scaled conditions, resulting in similar estimated vs actual Δamplitude values and a mean estimation error close to 0 (Fig. 3C).

**Figure 3.**
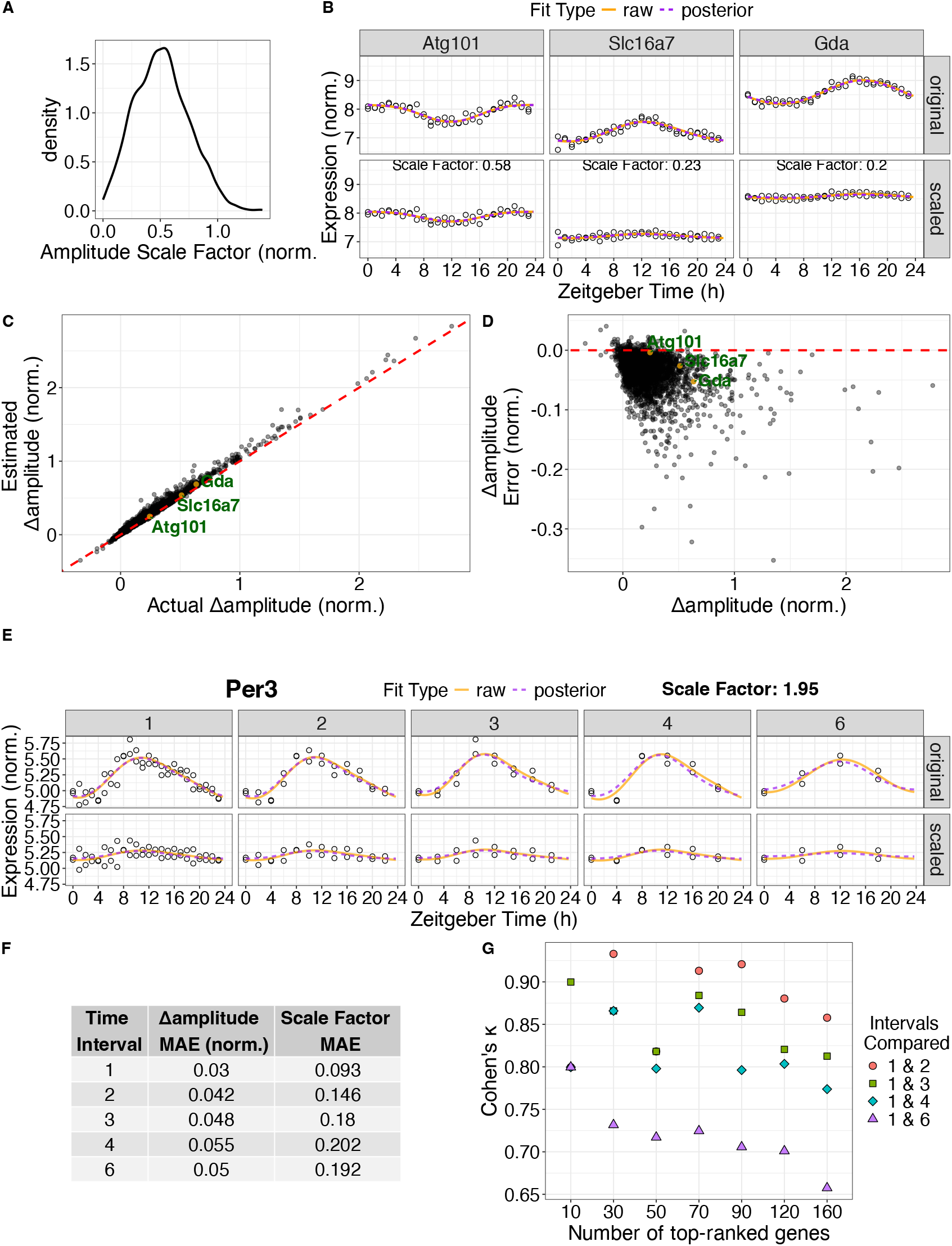
LimoRhyde2 estimation of artificial differential rhythmicity. **(A)** Density plot of generated scale factors (one value randomly assigned to each gene) applied to samples of all genes to create the ‘scaled’ condition for artificially derived differential rhythmicity. Dataset: GSE11923. **(B)** Time-courses of expression in the original (top row) and scaled (bottom row) conditions for select genes. Scale factors used per gene printed atop the scaled condition. Points represent samples. Curves represent LimoRhyde2 fitted curves of the labeled type. **(C)** Scatter plot of the calculated (actual) Δamplitude between original and scaled conditions vs the LimoRhyde2 estimated Δamplitude, with selected genes labeled in **(B)**. Points represent genes. Dashed lines indicate y = x. **(D)** Scatter plot of the estimation error (actual - estimated Δamplitude) vs. estimated Δamplitude. Selected genes labeled in **(B)**. The dashed red line indicates y = 0. (E) Time-courses of expression in the original (top row) and scaled (bottom row) conditions for gene *Per3* (with associated scale factor) after down-sampling data at different time intervals (leftmost column represents original sampling interval of 1 h). Points represent samples. Curves represent LimoRhyde2 fitted curves of the labeled type. **(F)** Table showing mean absolute error (MAE) of Δamplitude and scale factor after running differential rhythmicity analysis at different time intervals. **(G)** Inter-rater agreement, as quantified by Cohen’s *κ*, comparing top-ranked genes based on LimoRhyde2 Δamplitude sampled at 1 h time intervals and consecutively decreasing time intervals.

As there is a relationship between the original and the scaled condition via the scale factor (see equations in Methods), we can also determine accuracy by calculating the scale factor between LimoRhyde2 estimates of original and scaled gene amplitudes (termed the estimated scale factor). When compared to the randomly assigned scale factor (actual) LimoRhyde2 consistently estimated values close to the actual, resulting in a mean estimation error near 0 (Fig. S1A). Genes with higher ‘scaled’ amplitudes tended to exhibit slightly larger estimation errors compared to genes with smaller values. This may result from LimoRhyde2 under-weighting estimates for genes with scale factors much greater than or much less than 1, since the majority of genes had scale factors close to 1 (Fig. 3D, Fig. S1B-C). Notably, this disparity disappeared when a constant scale factor was applied to all genes (Fig. S1D).

As we performed differential rhythmicity at a sampling frequency seldom used during genomic analyses, we evaluated the method’s accuracy when sampling at larger time intervals. We conducted differential rhythmicity analysis between original and scaled conditions additionally at 2h, 3h, 4h, and 6h intervals by removing samples by multiples of the sampling interval over the 24h period. The accuracy of LimoRhyde2 values tended to decrease with longer intervals, possibly because larger time intervals produced lower signal-to-noise ratios as more samples were excluded. (Fig. 3E-F, Fig. S1E-F). However, genes tended to maintain similar ranks across sampling frequencies when we compared gene rankings based on amplitude at 1-hour resolution with those at all other intervals (Fig. 3G). This suggests that LimoRhyde2 may provide comparable interpretations of the strength of gene rhythms despite variations in sampling intervals. To further understand LimoRhyde2’s performance limitations, we tested the stability and reliability of estimated differential rhythm statistics in response to data variation and uncertainty. We ran bootstrap resampling on RNA-seq data in which (SCN) tissue was sampled every 4 h for 24 h, with three replicates per time point (GSE72095) (Pembroke et al., 2015). First, we used the method described above to create an ‘original’ and ‘scaled’ condition. During each bootstrap, we reconstructed each gene’s time course by randomly sampling one of the three replicates per time point in each condition and performed differential rhythmicity analysis with LimoRhyde2. We performed 500 bootstrap resamples. Genes were assigned the same scale factor across all bootstraps (Fig. 4A).

**Figure 4.**
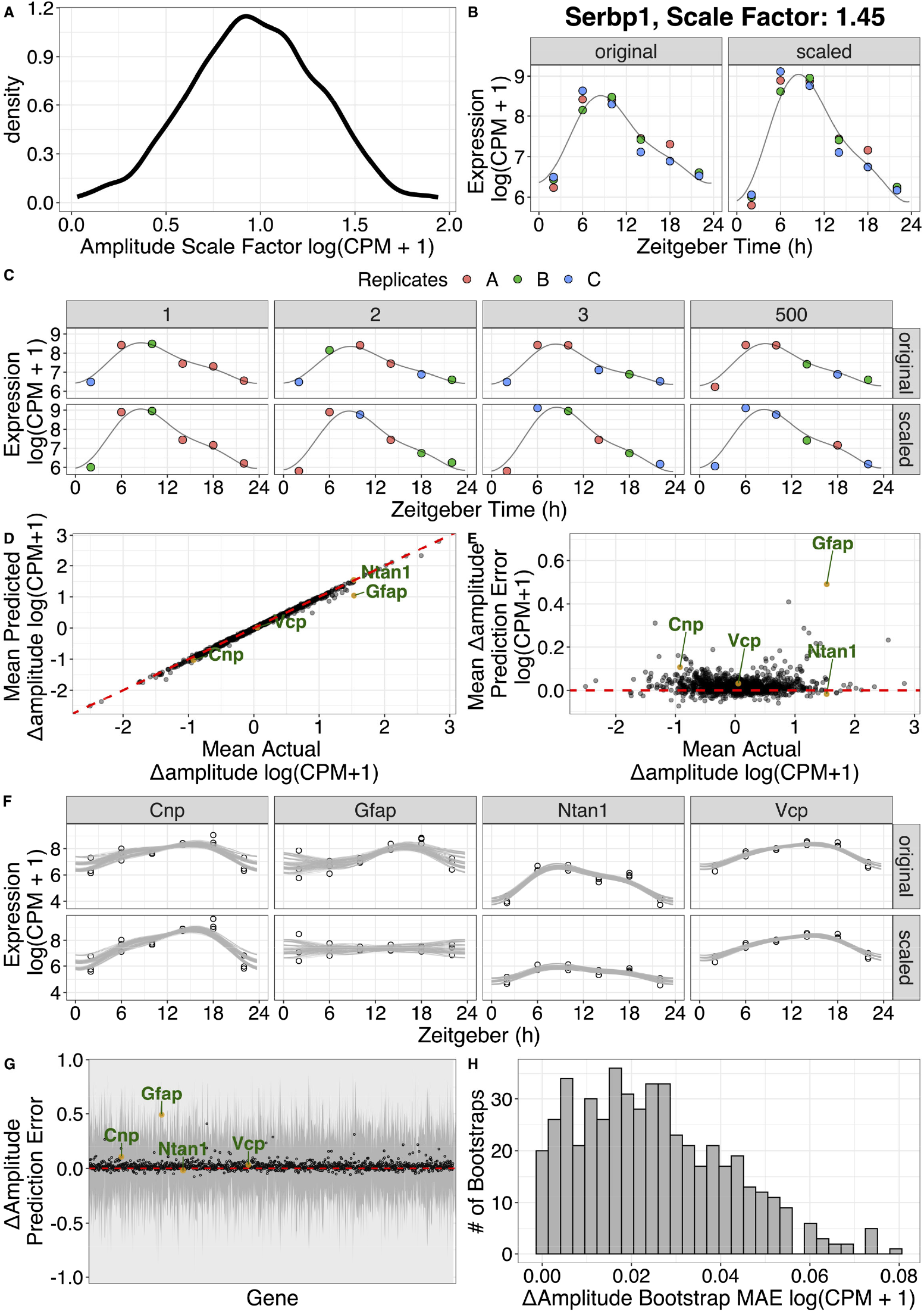
LimoRhyde2 differential rhythmicity accuracy from bootstrapping. **(A)** Density plot of generated scale factors (one value randomly assigned to each gene) applied to gene samples measurements to create the ‘scaled’ condition for artificial differential rhythmicity. Dataset used (GSE72095). **(B)** Time-courses of expression for gene *Serbp1* (with its assigned scale factor) in the original (left) and scaled (right) conditions. Points represent samples. Curves represent LimoRhyde2 posterior fitted curves. Color represents replicates. **(C)** Time-courses of expression in the original (top row) and scaled (bottom row) conditions for gene *Serbp1* at 4 different bootstrap resample runs of differential rhythmicity analysis with LimoRhyde2. Points represent samples. Curves represent LimoRhyde2 posterior fitted curves. Color represents replicate numbers. **(D)** Scatter plot of mean estimated Δamplitude and mean actual Δamplitude along with select genes (labeled in green). Points represent genes. Dashed lines indicate y = x. Values are averaged over 500 bootstrap runs. **(E)** Scatter plot of mean Δamplitude estimation error vs means actual Δamplitude, along with selected genes (labeled in green) Points represent genes. Dashed lines indicate y = 0. Values averaged across 500 bootstraps. **(F)** Time-courses of expression for genes labeled in C). Points represent samples. Lines represent fitted curves from 500 bootstraps. **(G)** Bootstrap Δamplitude estimation error and corresponding uncertainty. Points and gray ranges represent mean bootstrap prediction error and ± 1 s.d. per gene. **(H)** Histogram of bootstrap Δamplitude mean absolute error (MAE).

For each gene in a bootstrap event, we create a time course having 1 replicate per time point (Fig. 4B-C). Across all bootstraps, LimoRhyde2 consistently estimated Δamplitude and scale factors close to actual values, resulting in mean bootstrap estimation errors near 0 (Fig. 4D-E, Fig. S2A-B). As expected, LimoRhyde2 often produced conservative estimates of Δamplitude, leading to mean estimation errors consistently above 0. However, there was minimal correlation between the error in Δamplitude and its magnitude, with only a few genes having large errors at high Δamplitudes (Fig. S2B). In addition, genes having variable estimates across bootstraps tended to have noisier expression time courses, larger standard errors across bootstraps and showed similarly large LimoRhyde2-calculated credible intervals after sampling from the posterior distribution (Fig. 4F-G, Fig. S2B-F). Lastly, most genes had negligible differences in their estimation errors (Fig. 4H).

To additionally validate LimoRhyde2 using null differential rhythmicity, we divided the original Hughes et. al. (filtered for rhythmic genes) into two conditions by grouping samples from even-numbered and odd-numbered time points for each gene (Fig. S3). We then used LimoRhyde2 to quantify rhythmicity in each condition and differential rhythmicity between even and odd conditions. Given that this new data had half the number of samples as the original, we also performed rhythmicity analysis on the full course (referred to as the ‘original’ condition) to adequately analyze how sample dropout affects resulting estimates (Fig. 5A-B). As expected, LimoRhyde2 estimated similar amplitudes between even and odd conditions, resulting in near 0 Δamplitudes which did not correlate with the original amplitude (Fig. 5C, Fig. S4A). While amplitude values were similar between the two conditions, we saw differences in the density distributions between even, odd, and the original data, suggesting that there might be small but random changes in rhythmicity from dividing the data into two conditions (Fig. 5D).

**Figure 5.**
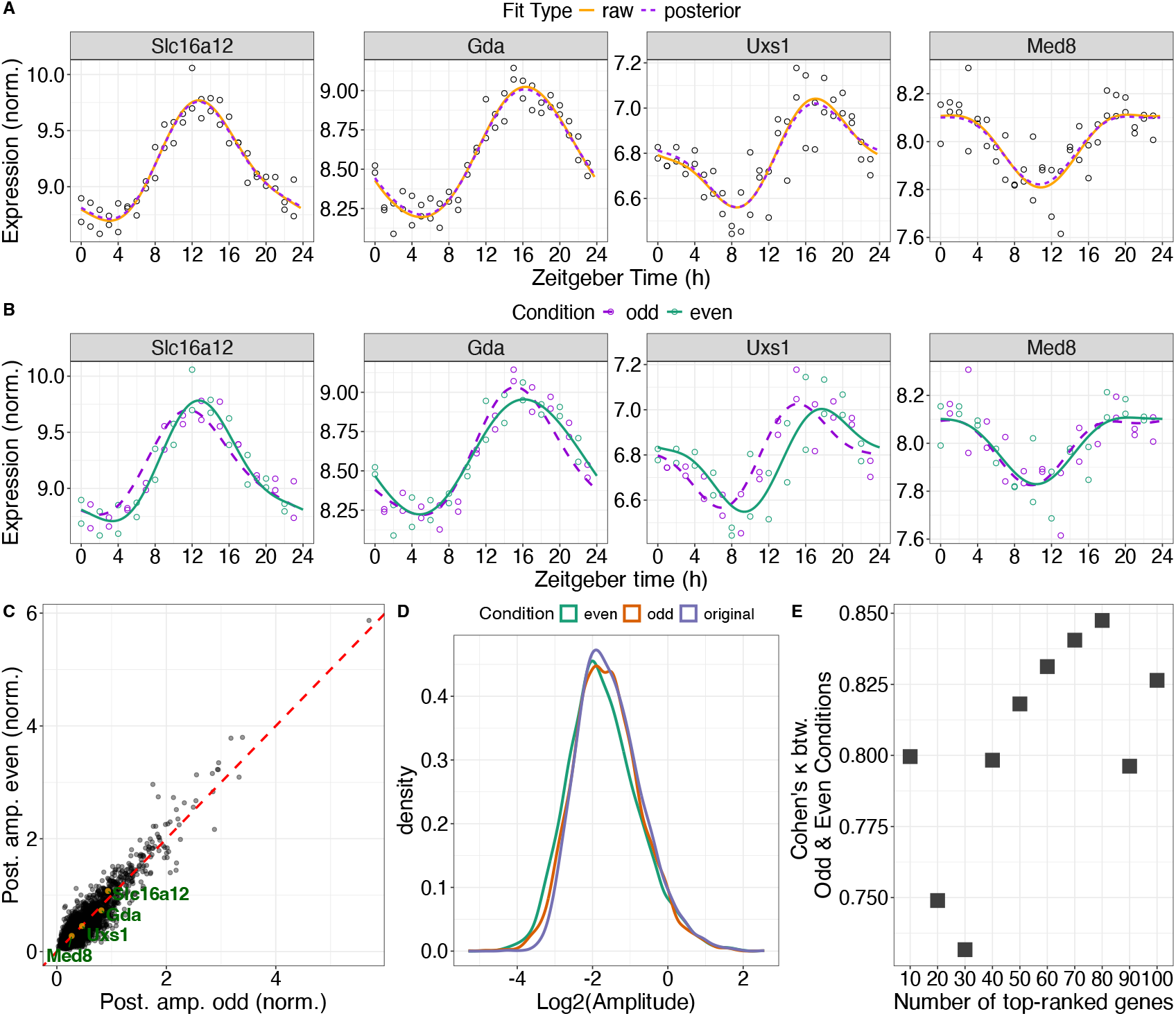
Quantifying null differential rhythmicity with LimoRhyde2. **(A)** Time-courses of expression for select genes. Points represent samples. Curves represent LimoRhyde2 fitted curves of the given type. Dataset used: (GSE11923). **(B)** Time-courses of expression for genes labeled in **(A)** with LimoRhyde2 fitted curves for odd (purple) and even (green) samples separately. Points represent samples. Color represents condition. Curves represent LimoRhyde2 posterior fitted curves. **(C)** Scatter plot of LimoRhyde2 amplitude calculated in the even condition vs the odd condition. **(D)** Density plots of log2-transformed amplitude. Color represents condition. Dashed lines indicate y = x. **(E)** Interrater agreement, as quantified by Cohen’s *κ*, comparing top-ranked genes based on LimoRhyde2 amplitude in odd and even conditions.

Perhaps due to increased standard errors in the new conditions compared to the original single one, LimoRhyde2 often calculated slightly different fitted curves for genes between conditions, likely due to the presence of within-condition trends in effects (Fig. 5B-D, Fig. S4B-C). Furthermore, genes with larger condition differences in standard error showed slightly larger differences in amplitude, even as changes occurred equally likely in each condition, supporting the random nature of this differential rhythmicity (Fig. S4D-F). Yet, LimoRhyde2 amplitudes led to similar top-ranked genes in both conditions, suggesting that random-generated condition-specific effects did not translate into different biological interpretations between conditions (Fig. 5E). Thus, LimoRhyde2 seems to preserve null differential rhythmicity, even as we expect some randomly occurring differential rhythmicity by artificially splitting the dataset into two.

## DISCUSSION

As circadian genomic experiments grow in complexity, statistical methods must grow with them to provide reliable means of interpreting such data. Rhythmicity parameters have long since provided a rich way of characterizing periodic variables (Cui et al., 2023, St. John et al., 2014, WILKINS et al., 2009) and their reliable quantification in genomic analyses may further elucidate richer details about the circadian system, especially about its response to circadian disruption and its degrees of effects in disease (Rijo-Ferreira and Takahashi, 2019). To enable more prescriptive analyses, circadian scientists must have tools that go beyond the mere identification of differential responses. For instance, Guan et al. analyzed differential rhythms by intricately tracking gene amplitude changes to genetic and environmental treatments to suggest the importance of cell-cell communication to circadian function within nearby cell types in the liver (Guan et al., 2020).

Outside of this and a few more studies, our understanding of the significance of rhythmic gene phase and to a larger extent amplitude in facilitating biological function remains limited. The endogenous circadian period is of biological origin and has clear relevance across various scales (Honma et al., 1987, Pagani et al., 2010). Some indications suggest that phase plays a biologically significant role in establishing systemic coordination, potentially serving to integrate specific functions (Evans et al., 2015, Kervezee et al., 2019, Welsh et al., 2004, Zhang et al., 2016). Our understanding of the potential role of amplitude is less comprehensive. Amplitude may be involved in shaping rhythmic biological processes, as diseases and other disruptions to the circadian clock can diminish amplitudes in rhythmic genes (Kohsaka et al., 2007, Teeple et al., 2023). Furthermore, the role of amplitude may offer insights into the intricate workings of the system, as compensatory mechanisms might exist to bolster amplitudes in genes when the clock is disrupted or when diseases induce disturbances (Crosby et al., 2019). Thus, further developing statistical methods to reliably quantify rhythmic parameters becomes imperative.

Here we present a well-validated process for extending LimoRhyde2 for differential rhythmicity analysis. LimoRhyde2 provides multiple outputs: estimated curves for each feature in each condition, effect sizes for various differential rhythm statistics, and credible intervals to evaluate uncertainty. While multiple outputs, as opposed to a single output (e.g., a *p*-value), may complicate the analysis process, the LimoRhyde2 framework offers a well-rounded interpretation for rhythmic features independent of strict yet arbitrary cutoffs. Lastly, LimoRhyde2’s effect distributions make it easier to incorporate differential rhythmicity results into follow-up analytical pipelines (Harrison et al., 2019). While this manuscript highlights how the method prioritizes important differentially rhythmic effects, it’s important to note that conceptually, differential rhythmicity can be equally beneficial as a distribution. This can help describe correlated patterns of effects, such as patterns of amplitude reprogramming observed between different tissues following a specific treatment Gal Manella (Manella et al., 2021). By prioritizing large effects in this manner, one can easily generate hypotheses about important targets by using biologically relevant parameters (i.e., amplitude and phase).

Obodo et al expounded on LimoRhyde2’s limitations in its initial paper. However, the method still has opportunities for future improvements in analyzing differential rhythmicity. Here, we focus primarily on validating LimoRhyde2’s accuracy in quantifying changes in amplitude. We largely assume the same high degrees of accuracy to apply when quantifying changes in phase. However, large-scale validation of phase using LimoRhyde2 should be followed to ensure accuracy. We have thus far only validated LimoRhyde2 on bulk transcriptome data, and are excited to apply it to other types of circadian genomic data, including data from ATAC-seq (Hor et al., 2019) and GRO-seq (Fang et al., 2021). Similarly, further validation is necessary to understand how LimoRyde2 can model differential rhythmicity in human genomic studies.

Studies use increasingly complex genome-scale experiments to address challenging questions in circadian biology as evidence of circadian genomic control continues to grow. Methods used to analyze rhythmicity from these experiments must accommodate the additional complexity to generate relevant conclusions. LimoRhyde2’s effect-size-based approach is a step in this direction. By directly estimating rhythmic properties that reflect circadian phenomena, LimoRhyde2 narrows the gap between statistical analysis and biological interpretation. For example, rhythmic properties can be compared between groups (i.e., cells, tissues, genotypes) in an experiment to ultimately assess clock coordination in whole organisms or to identify important time-dependent genes and potential drug targets. Additionally, by combining important analytical features (i.e., estimating curves, rhythmic properties, and uncertainty calculations) in one method, LimoRhyde2 can help standardize rhythmicity analyses and improve reproducibility. With these advances, LimoRhyde2 can help to expand investigations into biological rhythms and reveal new insights into the circadian systems.

## ACKNOWLEDGEMENTS

We thank Jake Hughey and Elliot Outland for their invaluable contributions and discussions regarding the ideas presented in this manuscript. Their expertise and feedback have greatly enhanced the depth of this work.

---

**Supplementary Figure 1:**
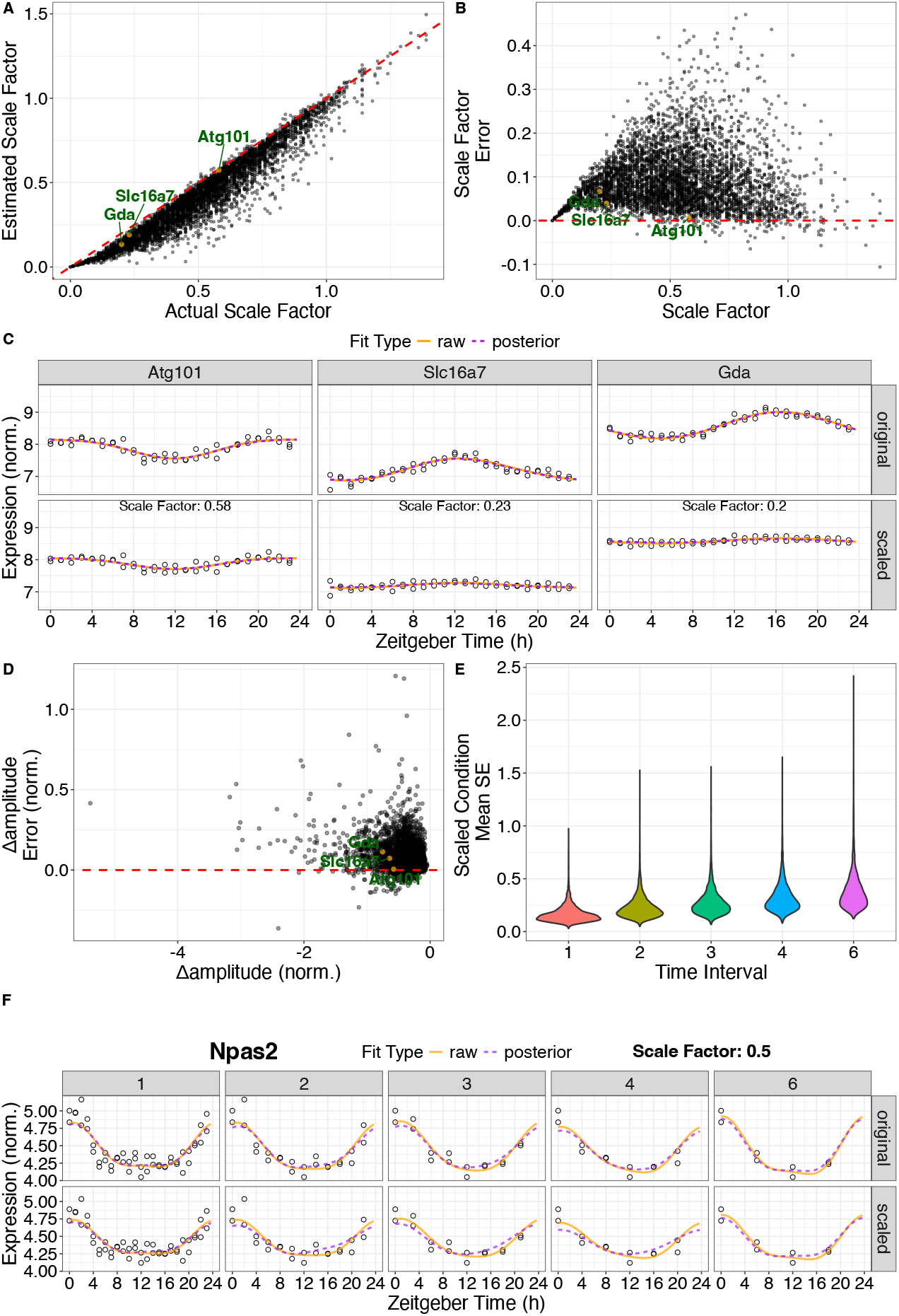
**(A)** Scatter plot of the actual vs estimated scale factor with 3 labeled genes. Points represent genes. Dashed lines indicate y = x. **(B)** Scatter plot of the scale factor estimation error (actual - estimated scale factor) vs. estimated scale factor along with genes labeled in **(A)**. Points represent genes. Dashed lines indicate y = 0. **(C)** Time courses of expression in the original (top row) and scaled (bottom row) conditions for select genes with scale factors per gene printed for the scaled condition. Points represent samples. Curves represent LimoRhyde2 fitted curves of the given type. **(D)** Scatter plot of the estimation error (actual - estimated Δamplitude) vs. estimated Δamplitude along with genes labeled in **(A)**. Values were calculated after performing differential rhythmicity analysis with LimoRhyde2 where the scaled condition was created with a constant scale factor applied to all genes. Points represent genes. Dashed lines indicate y = 0. **(E)** Violin plot showing mean standard error across genes in the scaled condition at five different time intervals. **(F)** Time-courses of expression in the original (top row) and scaled (bottom row) conditions for gene *Npas2* (with associated scale factor) after down-sampling data at different time intervals (leftmost column represents original sampling resolution of 1 h). Points represent samples. Curves represent LimoRhyde2 fitted curves of the given type.

**Supplementary Figure 2:**
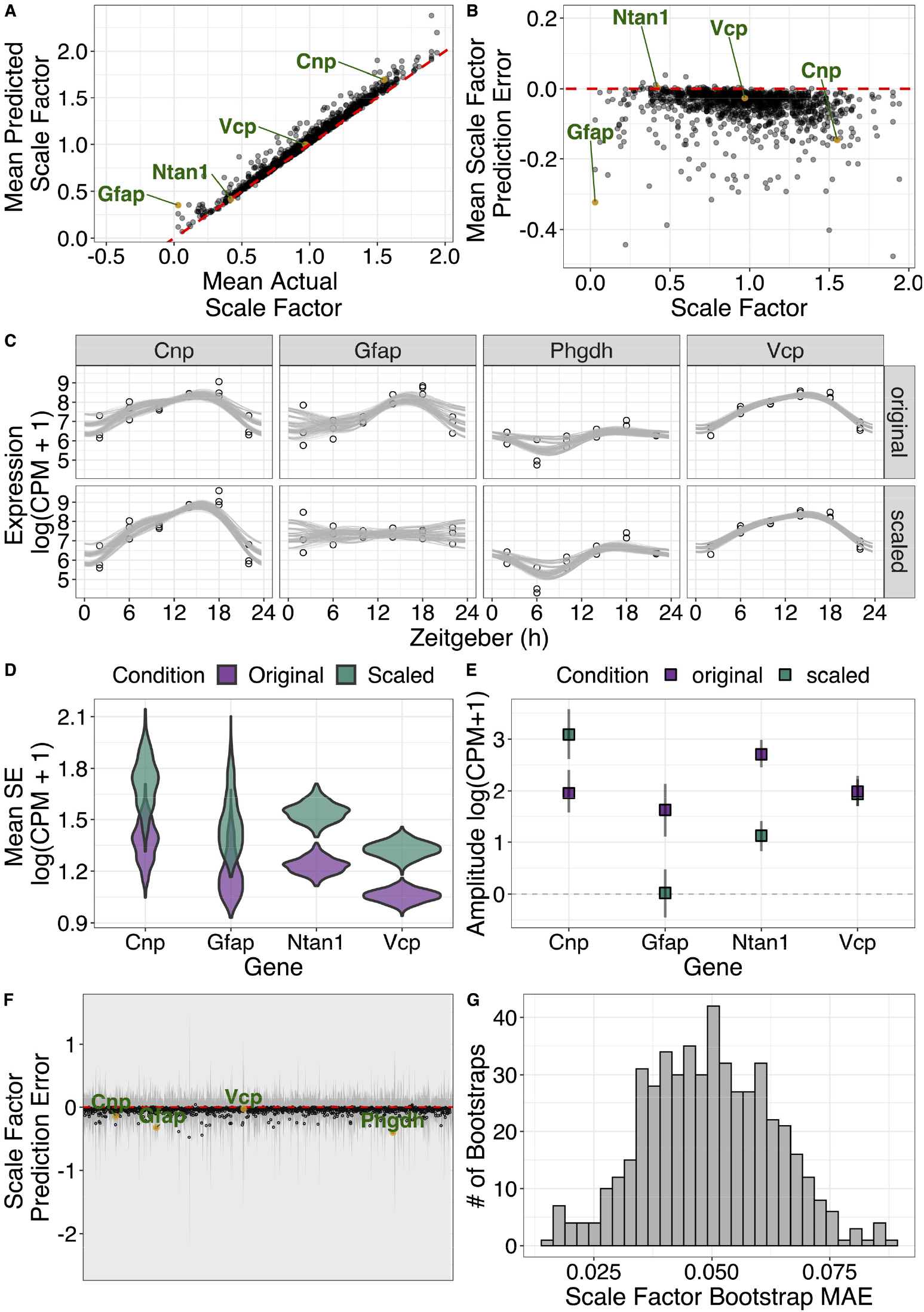
**(A)** Scatter plot of mean estimated scale factor and mean actual scale factor, with select genes. Points represent genes. Dashed lines indicate y = x. Values are averaged over 500 bootstraps using dataset GSE72095. **(B)** Scatter plot of mean scale factor estimation error vs mean actual scale factor, along with selected genes (labeled in green). Points represent genes. Dashed lines indicate y = 0. Values averaged across 500 bootstraps. **(C)** Time-courses of expression for genes labeled in **(A)**. Points represent samples. Lines represent fitted curves from 500 bootstraps. **(D)** Violin plot of mean standard error from bootstraps for selected genes in **(A)**. Color represents condition. **(E))** Posterior amplitudes and corresponding 90% credible intervals for selected genes in **(A)**. Points represent genes, color represents condition. Dashed line indicates 0 amplitude. **(F)** Point range plot of gene vs scale factor estimation error. Point and gray line represent estimation error mean and ± 1 s.d. of each gene. **(G)** Histogram of scale factor mean absolute errors.

**Supplementary Figure 3:**
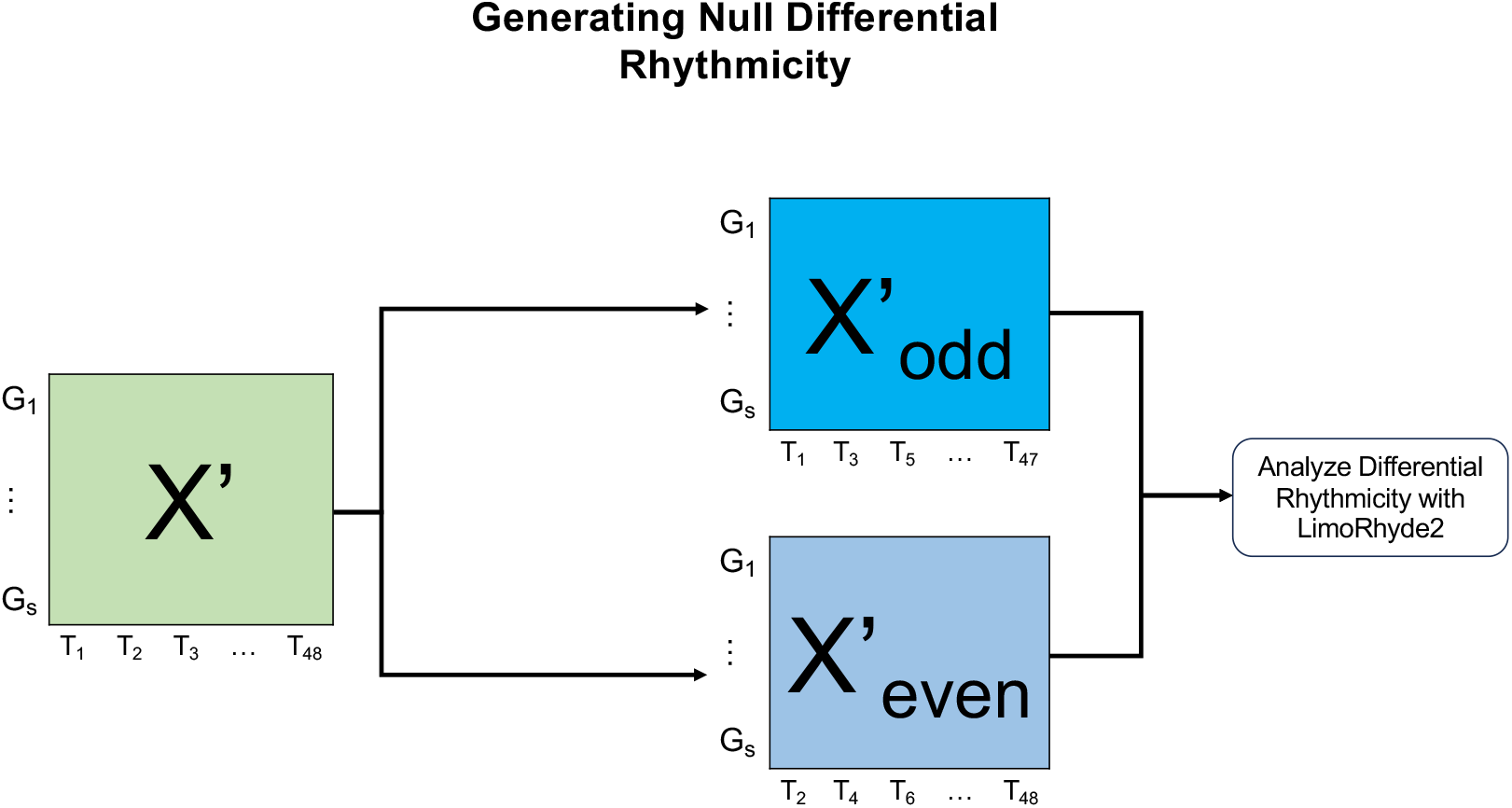
Generating null differential rhythmicity. Given a dataset X’ (filtered for identified rhythmic genes), with genes 1 to s and samples at time points T1 to T48, create two semi-equivalent conditions for LimoRhyde2 null differential rhythmicity analysis by diving samples occurring at odd (upper blue box) and even (lower blue box) time points. Analyze differential rhythmicity between odd and even conditions using LimoRhyde2

**Supplementary Figure 4:**
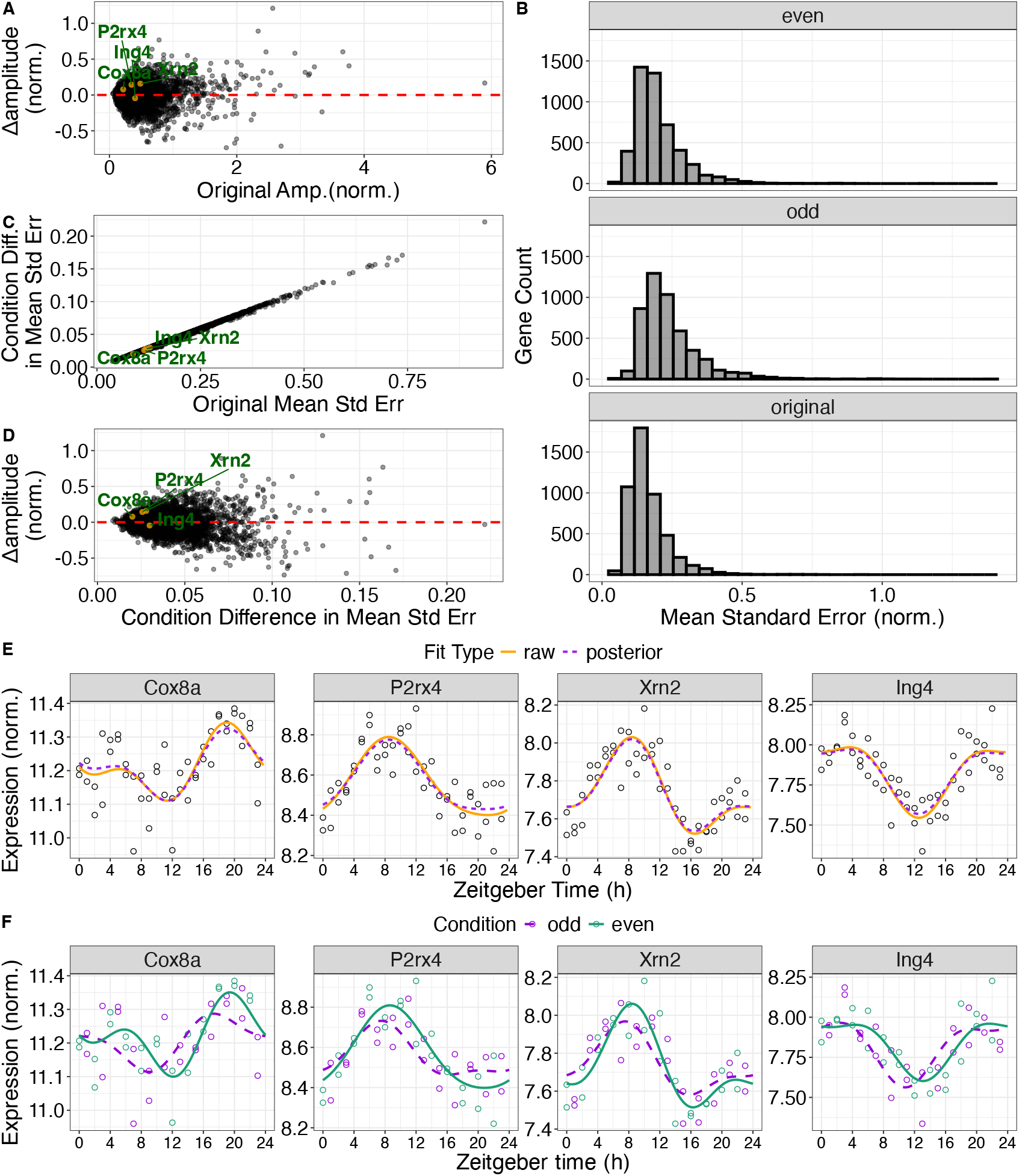
**(A)** Scatterplot of Δamplitude between even and odd conditions vs. amplitude from the original dataset, along with 3 selected genes. Points represent genes. Dashed lines indicate y = 0. Dataset used: (GSE11923) **(B)** Histogram showing counts of genes and mean standard error. Panels represent different conditions (and datasets). **(C)** Scatterplot of difference in mean standard error vs mean standard error in the original dataset with genes labeled in (A). Points represent genes. **(D)** Scatterplot of Δamplitude vs. condition difference in mean standard error. Points represent genes. Dashed lines indicate y = 0. **(E)** Time-courses of expression for select genes for their full-time course. Points represent samples. Curves represent LimoRhyde2 fitted curves of the given type. **(F)** Time-courses of expression for genes labeled in **(A)** with LimoRhyde2 fitted curves for odd and even samples separately. Points represent samples. Color represents condition. Curves represent LimoRhyde2 posterior fitted curves.

